# Age-Dependent Effects of Paraquat-Induced Parkinsonism on Serum Alpha-Synuclein Expression, Substantia Nigra Histoarchitecture and Neurobehaviour in Male Wistar Rats

**DOI:** 10.1101/2025.11.20.689379

**Authors:** Kelechi Emmanuel Ichie, Azuoma Lasbrey Asomugha, Izuchukwu Azuka Okafor

**Affiliations:** Department of Anatomy, Faculty of Basic Medical Sciences, College of Health Sciences, Nnamdi Azikiwe University, Nnewi Campus, PMB 5001, Nnewi, Anambra State, Nigeria; Division of Translational Anatomy, Department of Radiology, University of Massachusetts T.H., Chan School of Medicine, Worcester, MA, 01655, USA; Department of Medicine, Neurology Unit, Nnamdi Azikiwe University Teaching Hospital (NAUTH), Nnewi, Anambra State, Nigeria

**Keywords:** Paraquat, Parkinsonism, Alpha-synuclein, Substantia nigra, Age-dependence, Neurobehavior, Wistar rats

## Abstract

**Background:** Paraquat (PQ), a widely used herbicide, has been implicated in Parkinson’s disease (PD)-like neurodegeneration through oxidative stress and alpha-synuclein (α-syn) dysregulation. Age is a critical determinant of vulnerability to neurotoxins. This study investigated age-dependent effects of paraquat-induced Parkinsonism on serum α-syn expression, substantia nigra histoarchitecture, and neurobehavior in male Wistar rats.

**Methods:** Sixty-three male Wistar rats were stratified into juvenile, young-adult, and adult cohorts and assigned to control, paraquat, or PQ+Recovery groups (n = 7). Paraquat (10 mg/kg, i.p.) was administered twice weekly for three weeks; recovery animals were monitored for two months. Neurobehavioral tests including hanging wire, open field, and Y-maze were conducted at baseline, post-exposure, and post-recovery. Serum and substantia nigra α-syn were quantified using ELISA, and histology assessed cytoarchitectural changes. Data were analyzed using t-tests and ANOVA (p ≤ 0.05).

**Results:** Neurobehavioral outcomes showed no significant differences in spontaneous alternation across cohorts. Latency-to-fall remained stable in young-adult and juvenile rats but declined in adult PQ+Recovery animals (p = 0.05). Line crossings decreased in adult paraquat rats (p = 0.02). Urine pools decreased significantly in young-adult paraquat (p = 0.0010) and PQ+Recovery groups (p = 0.0016), with group effects in young-adult (p = 0.007) and juvenile cohorts (p = 0.047). Substantia nigra α-syn showed no significant differences (p > 0.22); serum α-syn differed in young adults (p = 0.039), with reduced levels in the recovery group (p = 0.047). Histology revealed paraquat-induced neuronal degeneration was most severe in adults, and partial structural restoration after recovery.

**Conclusion:** Paraquat induces mild, age-dependent neurotoxic effects, with adults showing the greatest vulnerability. Differential serum α-syn responses and histological findings highlight developmental stage as a key modulator of susceptibility and recovery following environmental neurotoxin exposure.

## Introduction

Parkinson disease (PD) is a chronic, progressive neurodegenerative disorder and the second most prevalent neurological disease, affecting more than 6 million people worldwide and approximately 1% of individuals over 60 years of age[1,2]. Clinically, PD is characterized by motor symptoms which include bradykinesia, rigidity, resting tremor, and postural instability, and a range of non-motor features including sleep disturbances, cognitive decline, depression, paresthesia, and autonomic dysfunction[3,4,5]. These motor disturbances primarily reflect selective degeneration of dopaminergic neurons in the substantia nigra pars compacta, which produces dopamine depletion in the striatum and impairs coordinated motor control[3].

Pathologically, PD is hallmarked by progressive nigrostriatal dopaminergic neuron loss and by accumulation of misfolded alpha-synuclein (α-syn) that aggregates into intracytoplasmic Lewy bodies[6,7,8]. Pre-synaptic α-syn which contributes to synaptic vesicle trafficking misfolds into oligomers and fibrils that disrupt neuronal integrity, induce oxidative stress and trigger neuroinflammation under pathological conditions[9]. Genetic alterations in the SNCA gene further promote α-syn aggregation and increase neuronal vulnerability in familial forms of PD[10]

Both biological and demographic factors, including sex and age, significantly influence PD risk and progression[11]. Epidemiological studies indicate higher incidence and greater clinical severity in males, possibly due to hormonal influences and elevated α-syn accumulation in male brains[12]. Estrogen appears to confer neuroprotection through antioxidant and anti-inflammatory mechanisms[13]. Aging, however, remains the most prominent risk factor, as it impairs mitochondrial function, diminishes proteasomal clearance, and elevates oxidative stress, thereby exacerbating α-syn aggregation and dopaminergic neuronal degeneration[14].

Both genetic susceptibility and environmental exposures contribute to Parkinson disease (PD) etiology[15,16]. Environmental factors, notably heavy metals, industrial chemicals, experimental toxins, herbicides and pesticides such as 6-hydroxydopamine (6-OHDA), MPTP, rotenone, and paraquat have been implicated in PD pathogenesis, with paraquat (1,1′-dimethyl-4,4′-bipyridinium dichloride) receiving particular attention owing to its robust epidemiological and experimental links to disease development[17,18]. Mechanistically, paraquat undergoes redox cycling to generate excessive reactive oxygen species (ROS), disrupt mitochondrial electron transport, and induce oxidative damage, mirroring pathological processes observed in idiopathic PD and supporting its use as a reliable neurotoxicant for modeling parkinsonism in animals[19,20].

Alpha-synuclein, long studied within the central nervous system, is increasingly detectable in peripheral biofluids—including cerebrospinal fluid (CSF), blood, plasma, and serum—opening a minimally invasive window onto Parkinson disease (PD) biology[21–23]. Although CSF remains informative, its invasive collection limits frequent or large-scale use, whereas serum and plasma offer accessible alternatives because peripheral blood cells (notably red blood cells and platelets) abundantly express α-syn and contribute to circulating levels[24,25]. Serum measurements capture both neuronal release and peripheral cellular contributions and can therefore reflect disease-related protein aggregation and clearance processes in living subjects[26,27]. Alterations in total, oligomeric, or phosphorylated forms of serum α-syn have been associated with PD pathogenesis and progression, supporting its biomarker potential[28–30]. This peripheral readout is especially pertinent in toxin-induced models (such as paraquat exposure) that can disrupt blood–brain barrier integrity and promote leakage of neuronal proteins into circulation, thereby linking central neuropathology with systemic manifestations[30]. Together, these features make serum α-syn a translationally relevant, ethically favorable tool for longitudinal monitoring, early detection, and comparative evaluation in studies of Parkinsonism.

Rodent models, particularly Wistar rats, are widely utilized to study paraquat-induced neurodegeneration due to their well-characterized neuroanatomy and behavioral responsiveness[31]. Following exposure, rats exhibit PD-like features such as dopaminergic neuronal loss in the SNpc, increased α-syn expression, and motor deficits[14]. Behavioral assessments provide functional correlates to neuropathological changes, enabling multidimensional model validation. Tests such as the rotarod[32], open field[33], and cylinder tests[34] measure locomotor coordination and motor asymmetry, while the pole and stepping tests assess bradykinesia[35–37]. Advanced gait systems, including CatWalk and DigiGait, quantify stride and stance alterations[38,39]. Cognitive and affective impairments are evaluated through paradigms such as the Y-maze, Novel Object Recognition, and sucrose preference tests[40–42].

Histologically, the substantia nigra displays densely packed dopaminergic neurons rich in neuromelanin within the pars compacta[43]. In PD, this architecture deteriorates, characterized by neuronal loss, reduced pigmentation, reactive gliosis, and accumulation of Lewy bodies[44]. Studying such histoarchitectural disruptions across different age cohorts provides critical insight into how aging and environmental stressors interact to drive neurodegeneration. Age-related increases in oxidative vulnerability intensify paraquat’s neurotoxic effects, reducing neuronal resilience and repair potential[45]. Thus, understanding these age-dependent trajectories is essential for developing interventions that address both the molecular and structural bases of PD pathology.

Although paraquat is well-established as a potent inducer of oxidative stress, dopaminergic neurodegeneration, and α-syn-related pathology on rodent models[20], critical gaps persist in understanding how these effects vary across the lifespan and whether peripheral biomarkers reliably reflect central injury. Most studies have focused on single age groups (typically adults)[46,47,48,49], despite compelling evidence that aging modulates mitochondrial function, proteostasis, and α-syn aggregation, all of which influence susceptibility to environmental neurotoxins[14]. Moreover, while α-syn mis-folding and aggregation are core features of PD[50], paraquat’s impact on total α-syn levels remains inconsistent, and few studies have explored whether serum α-syn mirrors central changes in toxin-induced parkinsonism. Recovery dynamics following paraquat exposure remain underexplored, yet evaluating post-exposure outcomes is essential for determining whether its neurotoxic effects are reversible and for identifying windows of neurorestorative potential[51]. As highlighted by recent reviews[31], inconsistencies in toxin-based PD models often stem from variability in age, strain, dosing, and exposure paradigms; factors that remain insufficiently addressed.

This study addresses major gaps in paraquat-based Parkinsonism research by comparing juvenile, young-adult, and adult male Wistar rats; quantifying α-syn expression in both serum and the substantia nigra; and evaluating neurobehavioral and histoarchitectural alterations alongside a two-month recovery phase. This integrative, age-stratified, and recovery-focused approach is designed to clarify how paraquat toxicity, α-syn dynamics, and neurorestorative processes evolve across developmental stages. Recovery assessment is particularly essential, as post-exposure models can reveal the brain’s intrinsic capacity for repair, compensatory mechanisms, and potential therapeutic windows[51].

This study is grounded in the hypothesis that paraquat exposure induces age-dependent neurobehavioral impairments and substantia nigra degeneration, accompanied by distinct alterations in serum and brain α-synuclein expression. Furthermore, it posits that younger organisms possess a greater capacity for functional and structural recovery following toxic insult. To test this hypothesis, this study investigates the age-dependent effects of paraquat-induced parkinsonism on serum α-syn expression and substantia nigra histoarchitecture in male Wistar rats, while examining recovery effects following exposure. By integrating biochemical, behavioral, and histological analyses, the study aims to elucidate age-related susceptibility to neurodegeneration, the interplay between environmental toxins and α-syn pathology, and the brain’s capacity for recovery—advancing the search for effective, age-tailored therapeutic strategies in Parkinson’s disease.

## MATERIALS AND METHODS

This study was conducted in the Department of Anatomy, College of Health Sciences, Nnamdi Azikiwe University, Nnewi Campus, Nigeria, over a period of three months (June–August 2025)

All experimental protocols were approved by the Animal Research Ethics Committee of Nnamdi Azikiwe University, Awka (Reference Number: NAU/AREC/2025/0059; issued 12th May 2025) and conducted in accordance with the National Institutes of Health (NIH) guidelines for the care and use of laboratory animals[52].

A total of 63 apparently healthy male Wistar rats were used for this study. The animals were obtained from a nursery animal farm at the College of Veterinary Medicine, Michael Okpara University of Agriculture, Umudike, Abia State. The animals were stratified into three age cohorts: Juvenile (6-8 weeks old), Young adult (>8–12 weeks old), and Adult (>12 weeks old) based on their age prior to any experimental exposure[53]. We acknowledge that over the course of the 13-week study, some juvenile animals transitioned into later developmental stages; however, categorization was maintained according to baseline age in order to preserve consistency of group assignment and experimental comparisons. In each cohort, animals were randomly grouped into three groups (n = 7 rats per group): control group received normal saline (10 mg/kg b.w.) intraperitoneally, the paraquat (PQ) group received paraquat dichloride (10 mg/kg b.w.) intraperitoneally twice weekly for three consecutive weeks, while the PQ+Recovery group received paraquat treatment paraquat dichloride (10 mg/kg b.w.) intraperitoneally twice weekly for three consecutive weeks, and were allowed a two-month recovery period post-exposure before sacrifice. Selection of a two-month recovery period for the study was based on the pharmacokinetics of paraquat in the brain. Research indicates that paraquat persists in the ventral midbrain of mice with a half-life of approximately one-month[54], suggesting significant retention over extended periods. This prolonged presence implies that a two-month period will potentially allow for substantial paraquat clearance, enabling the assessment of both immediate and long-term neurotoxic effects.

The animals were housed in standard polypropylene cages lined with wood shavings, under ambient conditions typically at 22 ± 2 °C and standard humidity, with an approximately 12-hour light/dark cycle. They had ad libitum access to standard laboratory rat chow and clean drinking water. Cages and bedding were changed thrice weekly. Animals were acclimatized for two weeks before the start of the experiment and health checks were conducted daily. [55].

### Induction of Parkinsonism

Paraquat dichloride (1,1′-dimethyl-4,4′-bipyridinium dichloride; analytical grade, Sigma-Aldrich, USA) stock solution with a concentration of 200 mg/mL was purchased for the study to induce parkinsonism. To prepare the working solution at a final concentration of 10 mg/mL, a 20-fold dilution was required. This was achieved by mixing one part of the stock solution with 19 parts of normal saline. Specifically, 1 mL of the 200 mg/mL paraquat stock was diluted with 19 mL of normal saline to yield 20 mL of a 10 mg/mL paraquat solution. Fresh dilutions were prepared each day of administration. Paraquat was administered intraperitoneally twice weekly over a period of three weeks, following the protocol described by Tariq et al. [56]. Prior to each administration, rats were weighed.Age-matched control rats received equivalent volumes of normal saline (administration vehicle).

The PQ+Recovery group was observed for two months after the final paraquat administration to evaluate potential long-term effects in neurobehavioral performance and alpha-synuclein expression. Mortality and the overall health status of the animals were carefully monitored throughout the study.

### Neurobehavioral Assessments

Behavioral testing was conducted before the administration of paraquat, 24 hours after the final paraquat injection in the experimental groups, and at the end of the recovery period for the recovery groups. All the tests were performed in a sound-attenuated environment during the daylight phase (09:00–14:00 hr), and all the apparatus used were cleaned with 70% ethanol between trials to eliminate olfactory cues.

The hanging wire test was performed to assess grip strength, neuromuscular endurance, and motor coordination. Each rat was placed on a horizontal metal wire (50 cm above the ground), and the latency to fall (in seconds) was recorded with a cut-off time of 240 seconds [*Figure 1(B)*]. Higher latencies indicated better coordination[57].

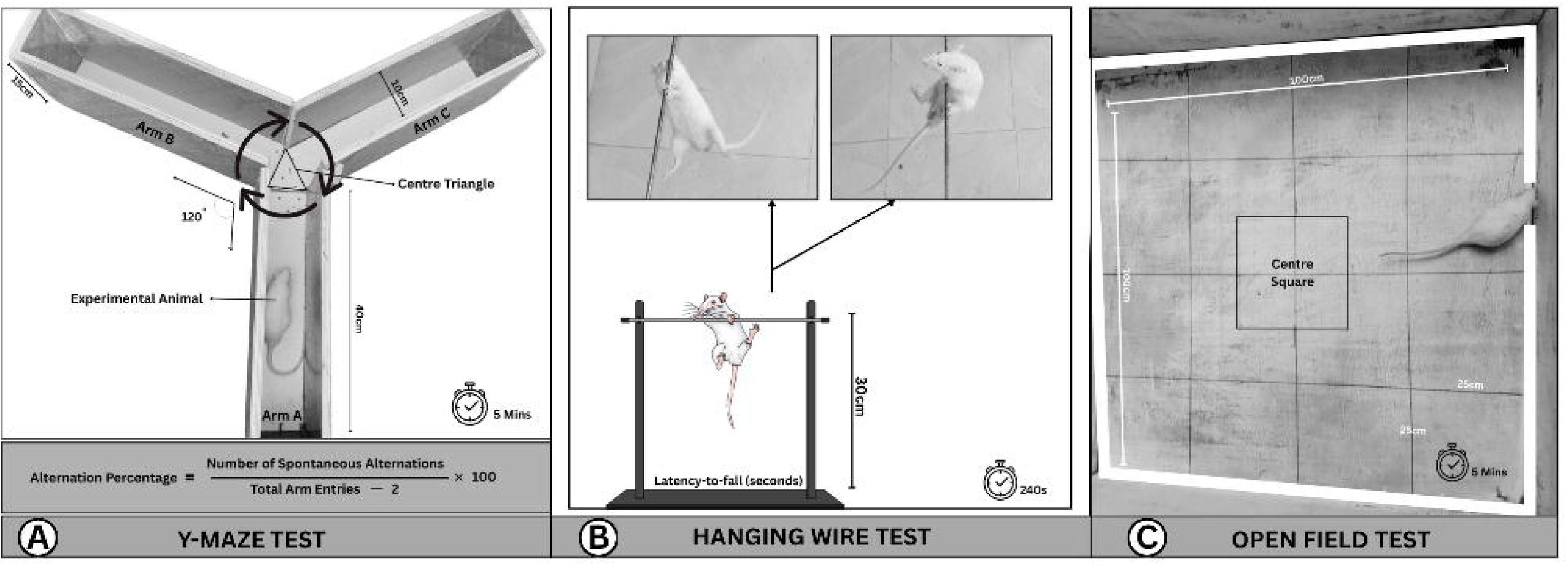

The Open Field Test (OFT) was used to assess locomotor activity and excretory behaviour. The apparatus consisted of a square arena (100 × 100 × 40 cm) with the floor divided into equal squares.

Rats were placed individually in the centre and allowed to explore for five minutes [*Figure 1(C)*]. Parameters recorded included line crossings (number of squares crossed with all four paws, reflecting locomotor activity), and number of urine pools (indicator of stress response). Higher locomotion and reduced excretory behaviour were interpreted as improved exploratory activity and anxiety-related behaviours respectively[58].

Spatial working memory was assessed using the Y-maze. The maze consisted of three arms (40 × 10 × 15 cm) arranged at 120° angles [*Figure 1(A)*]. Each rat was placed at the end of one arm and allowed to explore freely for five minutes. Spontaneous alternation behavior (SAB) performance was recorded when an animal had placed all 4 paws in the arm runway outside of the centre triangle. SAB performance was defined when an animal had entered each arm with 4 paws in a sequential order without returning to a previous arm (for example arm A to B followed by arm B to C). Higher alternation rates indicated improved spatial working memory[59].

### Animal Sacrifice

At the conclusion of experimental procedures, animals were euthanized by decapitation in accordance with institutional ethical guidelines and international standards for the care and use of laboratory animals. Prior to sacrifice, each rat was anaesthetized using intraperitoneal administration of ketamine (80 mg/kg) and xylazine (10 mg/kg). All procedures were performed by trained personnel under sterile conditions to minimize contamination and preserve tissue integrity.

### Blood Collection and Serum Preparation

Following confirmation of adequate anesthesia, blood was collected via ocular puncture. Blood samples were collected following the three weeks induction period (for control and paraquat groups), and after the two-month recovery period (for recovery groups) across all study cohorts. Samples were collected into plain tubes, allowed to clot at room temperature for 30 min, and centrifuged at 3000 rpm for 15 minutes at 4 °C. The serum was carefully separated, aliquoted, and stored at –20 °C until biochemical assays were conducted.

Following blood collection, the animals were subsequently euthanized by decapitation, in accordance with the AVMA Guidelines for the Euthanasia of Animals[60] and the approved Institutional Animal Care and Use Committee (IACUC) protocol, and the brains were rapidly excised and weighed. Relative weight of the brain was determined as organ weight divided by body weight. The substantia nigra of the left hemisphere was carefully dissected on ice, homogenized in phosphate-buffered saline (PBS, pH 7.4), and centrifuged at 10,000 rpm for 15 min at 4 °C. The supernatant was collected and stored at –80 °C until further biochemical analysis. Also, the substantia nigra of the right hemisphere was also carefully dissected and fixed in 10% neutral-buffered formalin for histological analysis.

### Tissue Processing and Light Microscopy

The substantia nigra of the right brain hemisphere were fixed in 10% neutral-buffered formalin at 4°C for 48 hours, dehydrated through graded alcohols, cleared in xylene, and embedded in paraffin. Serial coronal sections were cut on a rotary microtome at 5 µm thickness to provide complete cross-sections through the rostrocaudal extent of the hemisphere and mounted on glass slides. Sections were stained with hematoxylin and eosin (H&E) following standard procedures[61]. The substantia nigra pars compacta (SNpc) was identified on H&E-stained coronal sections by anatomical landmarks and characteristic histoarchitectural features. Representative sections spanning the SNpc were selected for analysis.

Photomicrographs were acquired from mounted H&E sections using a calibrated brightfield microscope (Labomed America Inc.; iVu 3100, LD400) connected to a PC (Dell Inc Latitide 3580, 1.4.1). Images were captured with the PixelPro application Version 8.0 on the PC using identical magnification (x400), illumination, and exposure settings for all samples to maintain consistency. Image acquisition and subsequent histological assessments were performed with the examiner being blinded to experimental groups. The photomicrographs were assessed for possible histological changes in the SNpc.

### Determination of Alpha-Synuclein Levels in the Serum and Substantia Nigra

Serum and substantia nigra alpha-synuclein concentration was quantified using the ELISA protocol with a sandwich ELISA kit obtained from Cell Signaling Technology, Inc., MA, USA. The kit manufacturer’s protocol was strictly followed for the α-syn estimation. Standards and samples were run in duplicates. The absorbance was read at 450 nm using a microplate reader (Bio-Rad, USA). Results were expressed as ng/mL of alpha-synuclein, and inter-assay coefficient of variation was kept below 10%.

### Statistical Analysis

All data were presented as mean ± standard deviation (SD). Within-group comparisons of pre- and post-neurobehavioral scores for each treatment group (control, paraquat, PQ+Recovery) in each age cohort (adult, young adult, juvenile) were evaluated with paired t-tests to determine the effect of paraquat within a specific group across the two time points. To assess the effect of treatment across age cohorts while accounting for the repeated pre/post measurements, a repeated-measures two-way ANOVA was applied, while one-way ANOVA was utilized for assessing effect of treatment within-cohorts.

All statistical analyses were performed using GraphPad Prism version 10.0 and IBM SPSS Statistics version 25.0 (IBM Corp., Armonk, NY, USA). Statistical significance was set at p ≤ 0.05, and exact p-values are reported for primary outcomes.

## RESULTS

### Relative Brain Weight

There was no significant difference in the relative brain weight of the control, paraquat-treated, and PQ+Recovery groups across the adult (p = 0.196, R² = 0.419) young adult (p = 0.168, R² = 0.449) and juvenile (p = 0.645, R² = 0.136) cohorts (*Table 1*).

**Table 1.**
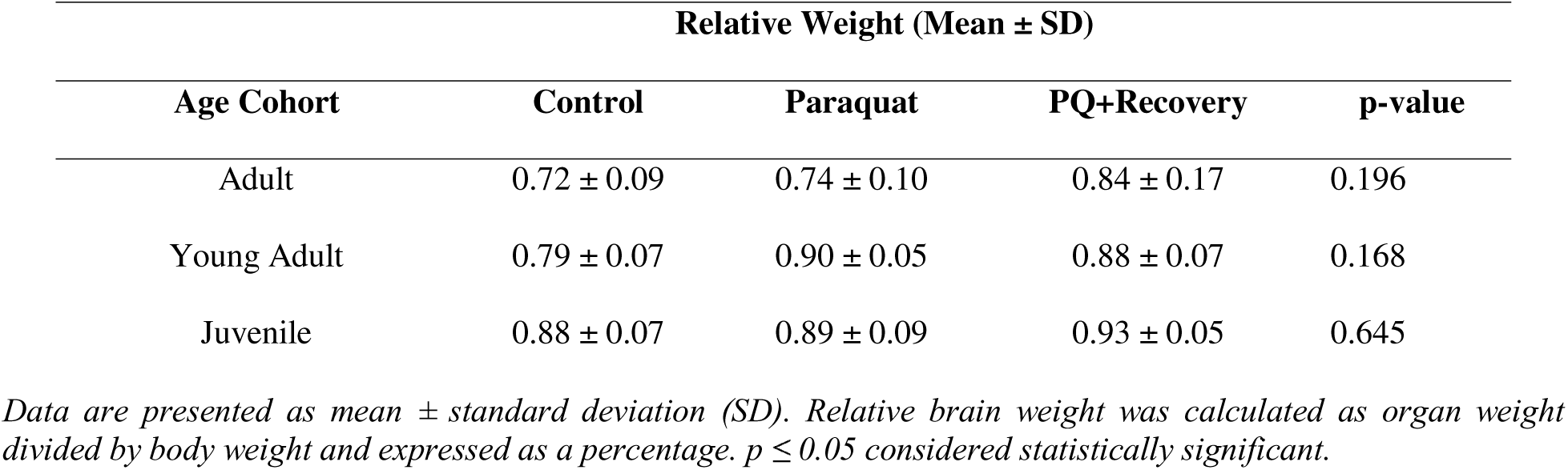
Relative brain weight across age cohorts following paraquat exposure and recovery.

### Neurobehavioural Assessments

#### Spontaneous Alternation (%)

There were no significant differences in pre- and post-spontaneous alternation test scores across experimental groups in all the age cohorts (p>0.05). There is no significant age-dose interaction across all groups in each age cohort (p>0.05). [*Figure 2A (i-iii)*]

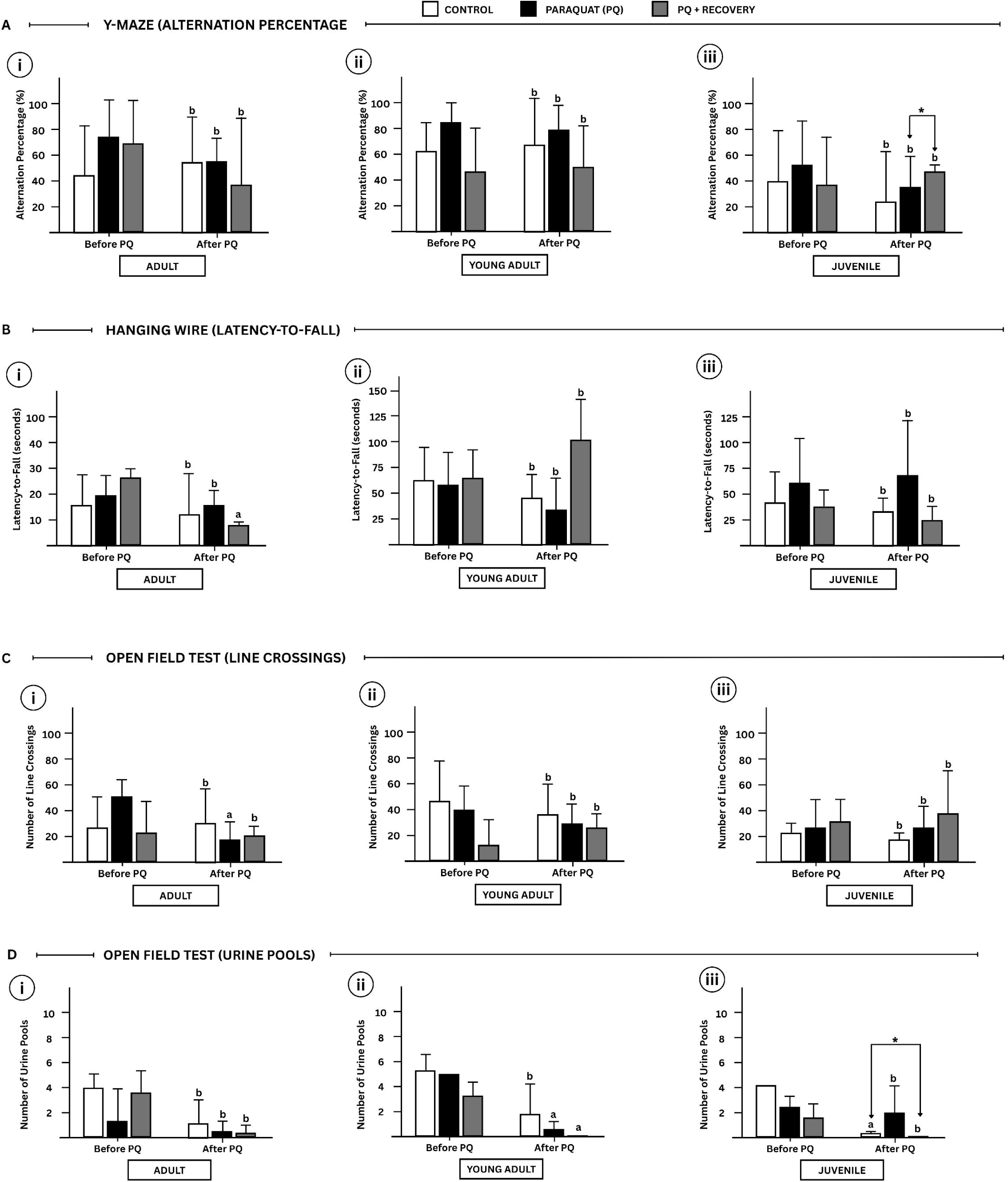

#### Latency-to-Fall (seconds)

Pre- and post latency-to-fall comparisons across experimental groups in all the age cohorts revealed no significant differences except in the PQ+recovery group of the adult cohort where a significant difference was detected (p =0.05). There is no significant age-dose interaction across all groups in each age cohort (p>0.05) [*Figure 2B (i-iii)*]

#### Number of Line Crossings (LC)

Pre- and post-LC test scores showed no significant differences among the experimental groups within each age cohort (p > 0.05), except in the adult paraquat group, which demonstrated a significant decline (p = 0.02). Additionally, no significant age–dose interaction was observed across any of the cohorts (p > 0.05).[*Figure 2C (i-iii)*]

#### Number of Urine Pools (UP)

In the pre- and post-UP test scores, there was no significant difference across the groups in the adult and juvenile cohorts (p>0.05); however, the paraquat group and the PQ+Recovery groups in the young-adult cohort were significantly different (p = 0.0010; p = 0.0016 respectively). Also, the PQ+Recovery showed a decrease in urination test scores in compared to the control group (p = 0.05). Additionally, no significant age–dose interaction was observed across the cohorts (p > 0.05). [*Figure 2D (i-iii)]*

### Alpha-Synuclein Concentration in the Substantia Nigra

The mean substantia nigra α-syn concentrations did not differ between control, paraquat, and paraquat+recovery groups in any age cohort (all p > 0.05). [*Figure 3(B)*].

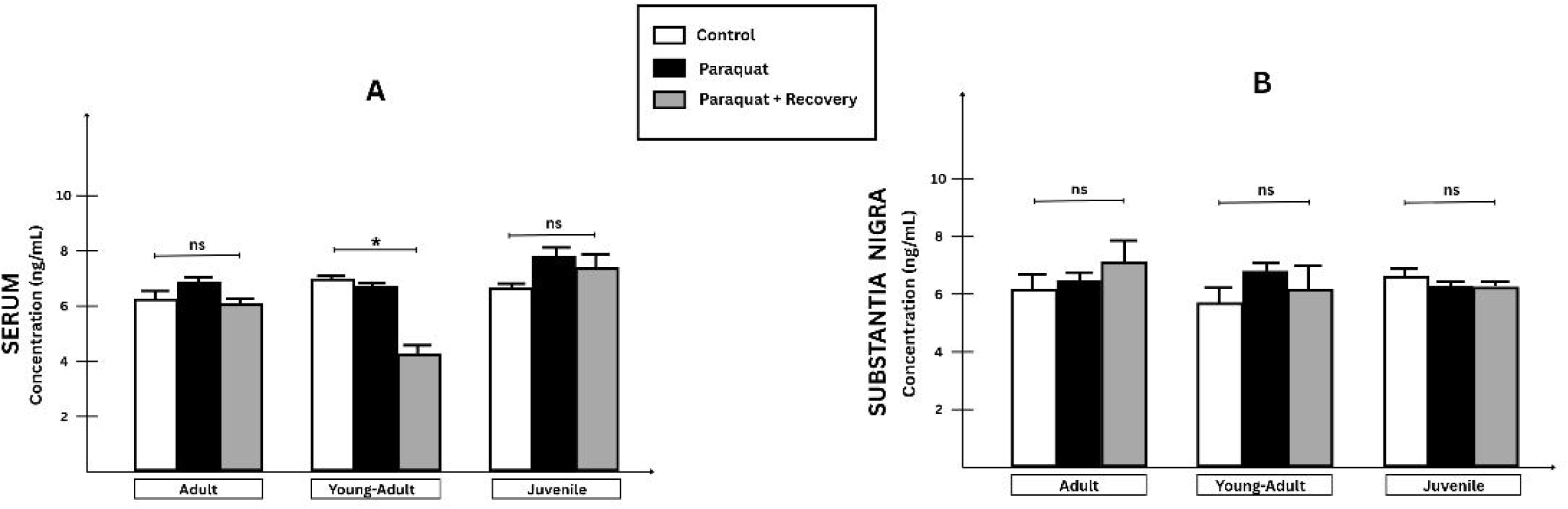

### Alpha-Synuclein Concentration in the Serum

The mean serum α-syn concentrations did not differ between control, paraquat, and paraquat+recovery groups in the adult and juvenile cohort (all p>0.05); however, in the young-adult cohort, there were significant group differences (p = 0.039). Post-hoc analysis further revealed that the recovery group had significantly lower alpha-synuclein concentrations compared to controls (p = 0.047). [*Figure 3(A)*].

### Histological Assessment of the Substantia Nigra

Histological examination of the substantia nigra pars compacta reveals distinct cytoarchitectural differences across experimental groups. In the control group across all cohorts, dopaminergic neurons exhibit well-preserved morphology (white arrow), and minimal evidence of vacuolation indicating structural integrity [*Figure 4; Plates a-c*].

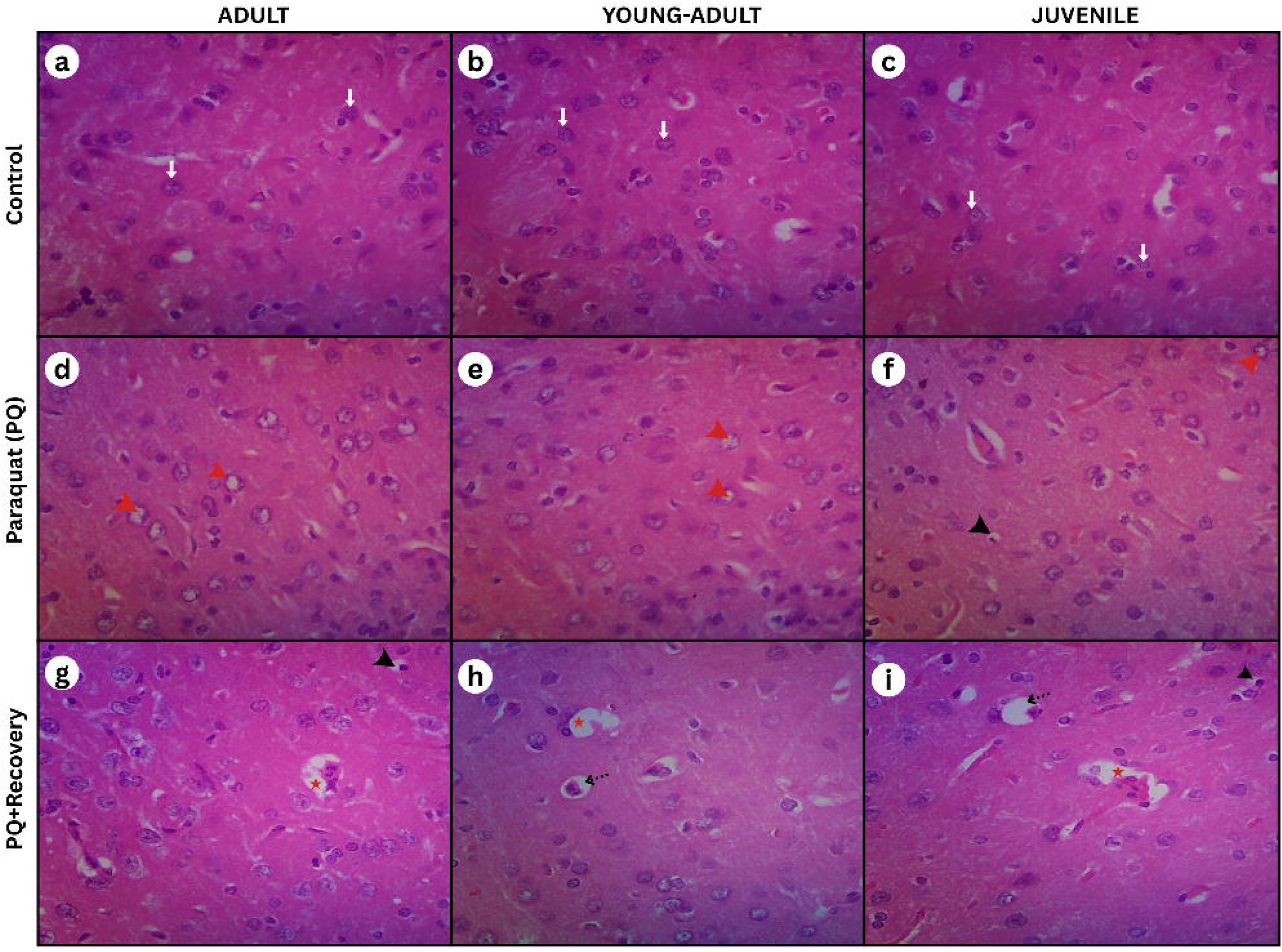

In contrast, paraquat-exposed groups show distinct histological alterations. Neurons, particularly in the adult cohort, display a pronounced “empty” cytoplasmic appearance (red arrowhead), suggestive of cytoplasmic shrinkage and loss of intracellular content. Similar degenerative changes and pyknotic nuclei (black arrowhead) are observed, though with progressively reduced severity in the young adult and juvenile cohorts. [*Figure 4; Plates d-f*].

In PQ+Recovery groups, cytoplasmic vacuolation (red star) and pericellular vacuolation (broken arrow) is evident with persistent pyknosis (black arrow head). However, there are observable improvements in neuronal density and cytoarchitecture. [*Figure 4; Plates g-i*]

## DISCUSSION

This study examined the effects of intra-peritoneal paraquat administration (10 mg/kg, twice weekly for three weeks) on neurobehavioral performance and α-syn expression in both the substantia nigra and serum of juvenile, young-adult, and adult Wistar rats. A two-month recovery phase was incorporated to assess the potential reversibility of paraquat-induced alterations. Our study demonstrates that intraperitoneal paraquat exposure (10 mg/kg, twice weekly for three weeks) produces age-dependent neurobehavioral and histological outcomes in male Wistar rats. Adult rats exhibited pronounced dopaminergic neuronal loss and degenerative changes in the substantia nigra, accompanied by motor deficits. In contrast, juvenile and young-adult rats showed only mild degeneration and largely preserved locomotor and cognitive function. Urination behavior revealed age-specific reductions in younger cohorts, suggesting subtle autonomic dysregulation that may precede overt neuronal loss. These findings indicate that both functional and structural impairments are closely linked and vary according to the maturity of the nigrostriatal system.

Our results largely align with previous studies demonstrating paraquat-induced motor deficits and nigrostriatal degeneration in rodents[62,46,49,63,64,65]. Fahim et al.[66] similarly reported reduced locomotor activity in Wistar rats following 10 mg/kg i.p. paraquat for three weeks, mirroring the adult cohort outcomes in our study. Conversely, Widdowson et al.[67] observed no deficits at lower oral doses, highlighting the importance of dose, route, and exposure frequency. Thiruchelvam et al.[68] also reported age-dependent susceptibility to paraquat and paraquatD+Dmaneb, with older animals showing progressive and irreversible motor and dopaminergic deficits, consistent with our findings.

Histologically, the pronounced nigral degeneration in adult rats supports the notion of age-related vulnerability, possibly driven by reduced antioxidative capacity and impaired mitochondrial resilience[69]. Our results corroborate prior work showing paraquat-induced dopaminergic degeneration[66,70,49,48,47,71]. However, studies such as Smeyne et al.[72] found no neuronal loss with similar dosing, likely due to differences in species, age, strain, or subtle methodological variations in administration and timing.

Our findings suggest that paraquat-induced deficits are both age- and severity-dependent. Adult dopaminergic neurons appear particularly vulnerable to oxidative stress and cytotoxicity[68], whereas younger neurons may tolerate early insult via compensatory mechanisms, as evidenced by stable spontaneous alternation scores and lack of motor deficits. Mild behavioral changes in juvenile and young-adult cohorts indicate sub-threshold damage that may progress with age, emphasizing the importance of early detection and longitudinal assessment. In humans, these results imply that older individuals or those with accelerated biological aging may be disproportionately susceptible to paraquat-induced dopaminergic injury. Following a two-month recovery period, partial restoration of substantia nigra pars compacta (SNpc) cytoarchitecture was observed. However, persistent pericellular vacuolation in older cohorts suggests sluggish structural recovery, consistent with the delayed regenerative capacity reported in older adults[73].

Relative brain weights did not differ significantly across treatment groups, suggesting that paraquat-induced neuronal damage at this dosage does not cause measurable brain atrophy. This suggests that neurodegeneration in this context is often microscopic rather than grossly morphological; hence, functional and cellular alterations may precede changes in gross anatomical indices[74], limiting the sensitivity of brain weight as a biomarker of neurotoxicity.

Mechanistically, paraquat triggers oxidative stress, mitochondrial impairment, and reactive oxygen species production, culminating in cytoplasmic shrinkage and nuclear pyknosis[46,65]. Genetic and strain-dependent factors may further influence susceptibility, as shown by Jiao et al.[75], who reported variable SNpc vulnerability across mouse strains. Such differences may similarly exist in Wistar rats and should be considered in interpreting paraquat toxicity outcomes.

### Alpha-Synuclein Expression in the Substantia Nigra and Serum

Paraquat exposure did not significantly elevate α-synuclein (α-syn) levels in the substantia nigra across age groups, aligning with prior observations that subacute or moderate paraquat dosing in wild-type rodents rarely increases total α-syn expression[76,77]. This reinforces the view that paraquat’s neurotoxicity is driven more by oxidative and mitochondrial stress than by increases in total α-syn. Indeed, paraquat preferentially induces α-syn misfolding, oligomerization, and conformational instability without affecting total protein abundance[78,45]. Upregulation of protective molecular chaperones such as HSP70[76] may further buffer against overt protein accumulation.

Serum α-syn showed age-dependent modulation, with young adults demonstrating notable reductions post-recovery, unlike juveniles and older adults. Such differences may reflect age-related variability in proteostasis efficiency, oxidative stress responses, and clearance pathways[79,80,30,28]. Because circulating α-syn originates largely from red blood cells and platelets[81–83], fluctuations in erythrocyte turnover, hemolysis, or platelet activation may influence serum concentrations.

The clinical literature provides an important parallel to these findings. Numerous ELISA-based studies have reported no significant difference in serum α-syn between Parkinson’s disease (PD) patients and controls[84–93]. A smaller group found lower serum α-syn in PD patients[94,95], whereas others documented higher levels[96–99]. This inconsistency mirrors our findings and reflects the strong influence of methodological variability, age, peripheral blood factors, and disease stage on α-syn measurements.

Non-ELISA assays—such as electroanalytical techniques[100], impedimetric assays[101], surface plasmon resonance [102,103], immunomagnetic reduction[104,105], and MSD electrochemiluminescence[106]—have similarly reported contradictory elevations or reductions. Collectively, these clinical data strengthen the interpretation that peripheral α-syn is a highly inconsistent biomarker, and that changes in its serum concentration do not reliably reflect central nigrostriatal pathology.

Thus, the lack of significant α-syn elevation in our paraquat-exposed groups aligns with broader preclinical and clinical evidence indicating that α-syn accumulation is not a universal feature of environmental toxicant models. Many toxin-based PD models, as highlighted by Dovonou et al. [31], fail to show robust α-syn pathology due to age, strain, and exposure-protocol differences.

Our findings reinforce a model in which paraquat induces reactive oxygen species (ROS) generation and NADPH oxidase activation[50,107], driving α-syn misfolding rather than overexpression. This suggests that future studies should quantify oligomeric, phosphorylated, or conformationally altered species using aggregate-specific assays. Age-stratified analyses of mitochondrial function, antioxidant capacity, and proteostasis markers would further clarify the relative resilience of younger rats. Given the variable nature of serum α-syn, complementary biomarker panels (e.g., NfL, DJ-1, inflammatory markers) and multi-omics approaches may improve translational relevance. Longitudinal tracking across extended recovery periods is also necessary to determine whether the mild alterations observed here progress with age or remain reversible.

